# SPE-51, a sperm secreted protein with an Immunoglobulin-like domain, is required for sperm-egg fusion in *C. elegans*

**DOI:** 10.1101/2021.07.07.451548

**Authors:** Xue Mei, Marina Druzhinina, Sunny Dharia, Amber R. Krauchunas, Julie Ni, Gunasekaran Singaravelu, Sam Guoping Gu, Diane C. Shakes, Barth D. Grant, Andrew W. Singson

## Abstract

Despite the importance of fertilization, the molecular basis of sperm-egg interaction is not well understood. In a forward genetics screen for fertility mutants in *Caenorhabditis elegans* we identified *spe-51*. Mutant worms make sperm that are unable to fertilize the oocyte but otherwise normal by all available measurements. The *spe-51* gene encodes a secreted protein that includes an immunoglobulin (Ig)-like domain and a hydrophobic sequence of amino acids. The SPE-51 protein acts cell-autonomously and localizes to the surface of the spermatozoa. This is the first example of a secreted protein required for the interactions between the sperm and egg with genetic validation for a specific function in fertilization. Our analyses of these genes begin to build a paradigm for sperm-secreted or reproductive tract-secreted proteins that coat the sperm surface and influence their survival, motility, and/or the ability to fertilize the egg.

## Introduction

Fertilization is a fundamental process in sexual reproduction during which gametes fuse to combine their genetic material and start the next generation in their life cycle. Fertilization involves species-specific recognition, adhesion, and fusion between the gametes (Bhakta et al., 2019; Schultz and Williams, 2005). In mammals, only a few proteins are known to be required for gamete interactions and have been validated with loss-of-function fertility phenotypes (Bhakta et al., 2019; Bianchi et al., 2014; Krauchunas et al., 2016; Mei and Singson, 2021). The mammalian egg coat, called the zona pellucida, contains ZP proteins among which ZP2 supports initial gamete recognition (Avella et al., 2016). After the sperm penetrates the zona pellucida, proteins on the plasma membranes of the sperm and egg mediate species-specific adhesion. To date, the only known binding receptor-pair is JUNO and IZUMO1 (Bianchi et al., 2014; Inoue et al., 2005). JUNO is a GPI-anchored protein on the egg surface; IZUMO1 is a single-pass transmembrane molecule with an Immunoglobulin (Ig)-like domain on the sperm. Crystal structures of the IZUMO1-JUNO complex suggest it is unlikely that this pair has fusogenic activity, but rather mediates adhesion between the two gametes (Aydin et al., 2016; Nishimura et al., 2016; Ohto et al., 2016). Several other molecules are also required for adhesion, including the Ig domain-containing SPACA6, TMEM95, SOF1, and FIMP in the sperm (Barbaux et al., 2020; Fujihara et al., 2020; Lamas-Toranzo et al., 2020; Lorenzetti et al., 2014; Noda et al., 2020). However, the binding partners for these proteins, if any, and their biochemical functions are unknown. The oocyte tetraspanin CD9 is involved in gamete interactions and/or egg surface membrane organization required for fertilization (Inoue et al., 2020; Kaji et al., 2000; Le Naour et al., 2000; Miyado et al., 2000). Identifying the molecules at the interface between the gametes has been a big challenge, because genetically manipulating these molecules leads to infertility and a genetic dead end (Bianchi and Wright, 2014; Krauchunas et al., 2016; Krauchunas and Singson, 2016; Mei and Singson, 2021). Further, defects in any of the processes that proceed direct gamete interactions, can lead to sterility thus complicating the classification of these proteins (Myles and Primakoff 1997).

The nematode *C. elegans* has been a powerful model to study the molecular basis of fertilization (Geldziler et al., 2011; Krauchunas et al., 2016; Marcello et al., 2013; Mei and Singson, 2021; Serrano-Saiz et al., 2019; Singson, 2001). Forward and reverse genetic approaches in *C. elegans* have identified a class of molecules that are required for sperm-egg interactions at the sperm plasma membrane (Krauchunas et al., 2016; L’Hernault et al., 1988; Nishimura and L’Hernault, 2010; Singaravelu et al., 2015; Singson et al., 1998). Mutations in this class of genes cause a unique Spe (spermatogenesis-defective) phenotype that is similar to that of the *Izumo1* mutant mice: mutant sperm show normal morphology and normal functions by all available measures but fail to fertilize the oocytes. To date, there are 8 previously described genes in this class: *spe-9*, *spe-13, fer-14, spe-38*, *spe-41*, *spe-42*, *spe-45*, and *spe-49* (Mei and Singson, 2021). These genes together are referred to as the *spe-9* class genes after the first gene of the group to be molecularly characterized (Singson, 2001; Singson et al., 1998). All these genes encode sperm surface transmembrane molecules that are thought to mediate the recognition, adhesion and/or signaling between the sperm and egg (Krauchunas et al., 2016; Marcello et al., 2013; Nishimura and L’Hernault, 2010). The non-redundant roles of these molecules suggest that they function with one another, and we refer to this functional unit as the fertilization synapse, similar to the neuronal or immune synapse (Krauchunas et al., 2016).

Here, we report the identification of another member of the *spe-9* class of genes, *spe-51*. The mutant phenotype of *spe-51* is highly similar to that of other *spe-9* class mutants. The SPE-51 protein also has a single Ig-like domain and a hydrophobic region, suggesting some exciting potential molecular functions in sperm-oocyte interactions. The protein acts cell-autonomously, and consistently, it localizes to the plasma membrane of the spermatozoa. Unexpectedly, instead of being a transmembrane protein like all the previously reported SPE-9 class proteins, SPE-51 is secreted. In another paper, we describe another secreted protein required for sperm function, SPE-36 (see Krauchunas, et al. on SPE-36 on Biorxiv). To the best of our knowledge, this is the first time that a secreted sperm molecule has been genetically demonstrated to be required cell autonomously for sperm-egg interactions at fertilization.

## Results

### *spe-51* mutants are sterile with a sperm specific fertility defect

Mutant *spe-51(as39)* hermaphrodites lay unfertilized eggs, occasionally producing one or two progeny (Figure 1). The *as39* allele is non-conditional and recessive. Homozygous hermaphrodites are sterile at all culture temperatures (Fig. 1A) and heterozygous animals are fully fertile (Fig. 1B). When mutant hermaphrodites are mated with wild-type males, they can produce progeny (406.2±12.0). To evaluate male fertility, we mated mutant *spe-51(as39)* males with *fem-1(hc17)* hermaphrodites that cannot produce sperm. We saw that while control males produce ∼500 progeny, *spe-51(as39)* males only produce 5.9(±1.2) progeny (Fig. 1C). Similar results were obtained when we mated mutant males with *dpy-5* hermaphrodites and counted outcross progeny (Fig. 2J). Thus, the mutant males are also sterile. These observations suggest that *spe-51* is required for fertility in both hermaphrodites and males.

**Figure 1.**
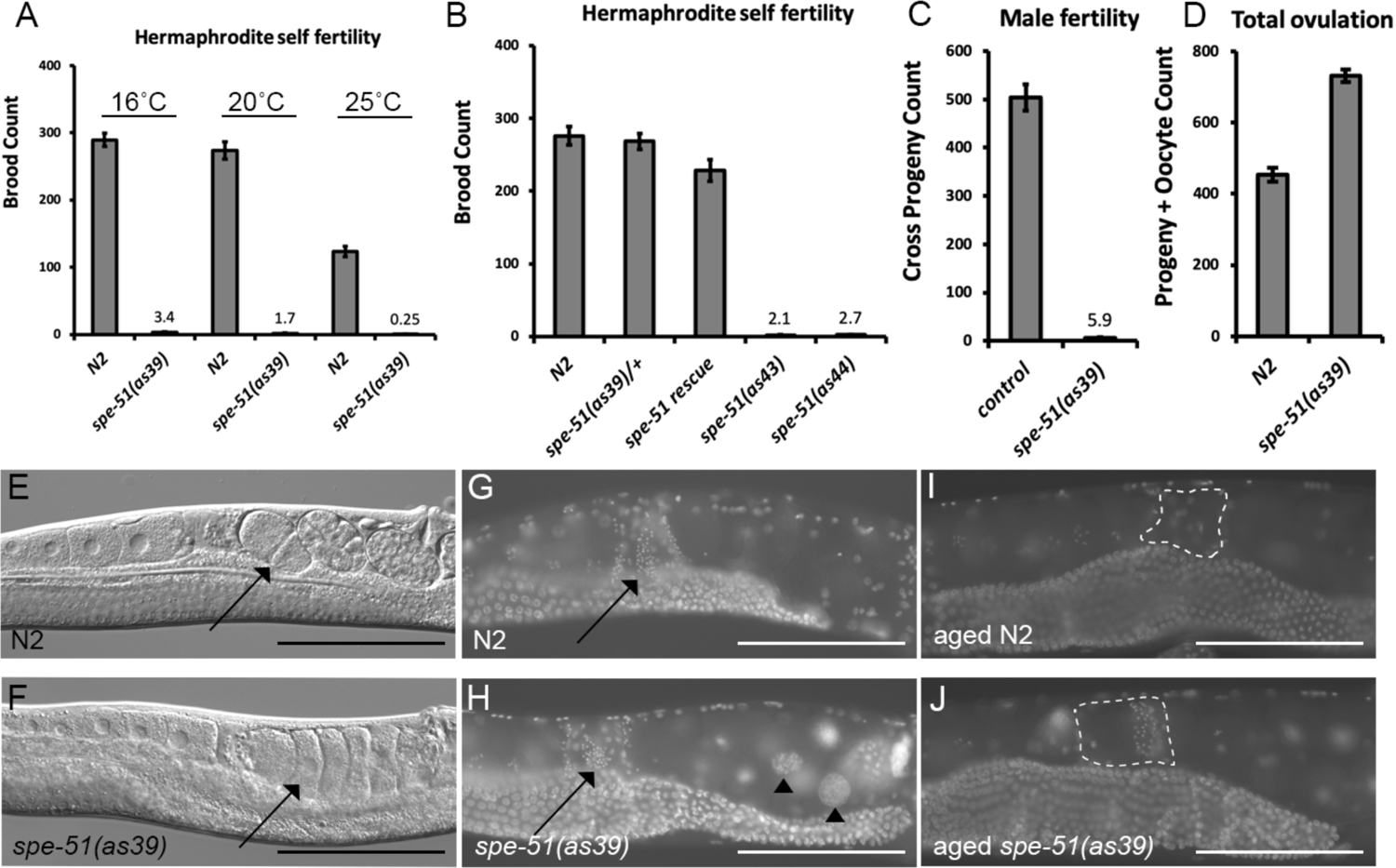
*spe-51(as39)* phenotypes. A. Progeny counts for N2 and *spe-51(as39)* hermaphrodites at different culture temperatures. B. Progeny count at 20 °C. C. Male fertility as measured by crossing mutant *spe-51(as39)* males with *fem-1(hc17)* hermaphrodites. D. Numbers of all the progeny and oocytes throughout the whole laying period. E-F, DIC imaging of the worm. Black arrows point to embryos or unfertilized oocytes in the uterus. G-J. DAPI staining of hermaphrodites. G-H, young adults; I-J, 4 days old adults. Black arrows in G and H point to sperm in the spermatheca. Black arrowheads in H point to endomitotic nuclei in the mutant. White dashed circles show the spermathecae in I and J. For A-D, Data are presented as mean ± SEM.

**Figure 2.**
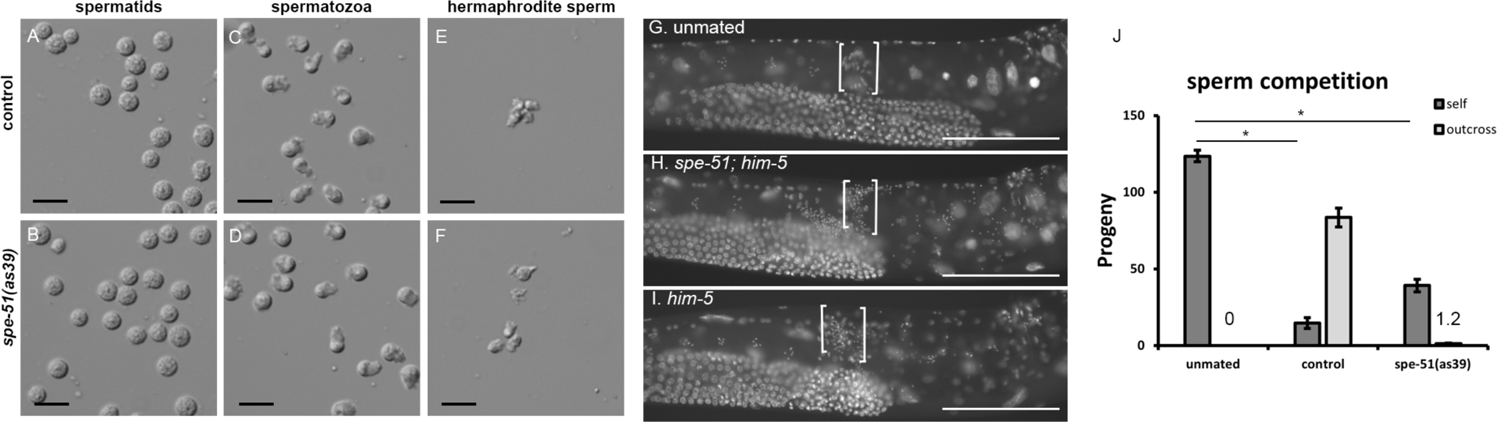
Sperm development in *spe-51(as39)* mutants. Spermatids are dissected from males. Spermatozoa are activated *in vitro* by Pronase. E and F, self-sperm from hermaphrodites. Scale bars in A-F represent 10 μm. G-I. DAPI stained *fem-1(hc17)* hermaphrodites. White brackets show the location of the spermathecae. Scale bars represent 50 μm. J, sperm competition. Progeny count of mated and unmated *dpy-5(e61)* hermaphrodites with control males or *spe-51(as39)* males. Data are presented as mean ± SEM. The number of self-progeny (Dpy) are significantly lower in hermaphrodites mated with either the control or *spe-51(as39)* males compared to unmated worms.

Adult worms of both sexes displayed no other mutant phenotypes than sperm specific infertility. We used light microscopy to further observe the defects in *spe-51(as39)*. We saw that the uteri of mutant hermaphrodites are filled with unshelled eggs (Fig. 1E-F, arrow) that had contacted and passed sperm in the spermatheca.

The morphology of the gonad looks normal in the mutant of both sexes. Maturing oocytes in the mutant also look indistinguishable from wild type. By DAPI staining, we found sperm are present in the spermatheca of the mutant (Fig. 1G-H). The unshelled eggs produced by unmated *spe-51* hermaphrodites showed a single DNA mass in the nucleus, the classic “endomitotic” phenotype that results from continuous DNA replication in the oocytes without chromosome segregation (Geldziler et al., 2011). Because chromosome segregation depends on centrosomes brought in by the sperm, our observation suggests that the sperm fails to enter the oocyte in *spe-51* mutants and those unshelled eggs are unfertilized.

In *C. elegans*, hormonal signals secreted by the sperm stimulate oocyte maturation and ovulation (Miller et al., 2001). Because we observed the presence of sperm in *spe-51(as39)* mutants, we asked whether ovulation rates were normal. We measured ovulation rates by counting the total number of unfertilized eggs plus progeny produced throughout the egg laying period. *spe-51(as39)* showed slightly higher ovulation totals than wild-type controls (Fig. 1D, t-test, p<0.001). Consistent with this observation, 4-day old mutants still have sperm in their spermathecae, as opposed to wild-type worms that have used up all the sperm by this time (Fig. 1I-J). We reason that the sperm in the mutant are not utilized for fertilization but retained for longer, and so induce more ovulation events. These data further suggest that the sterility in *spe-51* mutants is due to defects in the sperm that prevent them from fertilizing the oocytes.

### *spe-51* is not required for sperm activation, sperm motility or sperm competition

Since mutant *spe-51* sperm cannot fertilize the oocytes, we examined the morphology of mutant spermatids and spermatozoa. Spermatids are produced during meiosis and differentiate into spermatozoa after meiosis. In *C. elegans*, these round and non-polar spermatids differentiate into ameboid spermatozoa that are polarized and motile and use their pseudopods to crawl (Singson, 2001). This process, called sperm activation or spermiogenesis, can be induced *in vitro* by pharmacological reagents such as Pronase (Geldziler et al., 2011; Shakes and Ward, 1989). We initially observed the presence of sperm in mutant hermaphrodites, suggesting that spermatids were made (Fig. 1E-H). Indeed, we found plenty of spermatids when we dissected mutant males (Fig. 2A-B). These mutant spermatids appeared normal: they are round and have centered nuclei, indistinguishable from the wild type. We then asked whether these spermatids could activate *in vitro*. As shown in Fig. 2C-D, *spe-51(as39)* spermatids activated normally, compared to the wild-type controls. Consistently, mutant sperm from hermaphrodites were also activated, similar to those from the wild type (Fig 2E-F). Thus, *spe-51* is not required for spermatid production or sperm activation either *in vitro* or *in vivo*.

Sperm motility is critical for its function because the sperm needs the ability to crawl to the spermatheca, the site of fertilization in hermaphrodites (Geldziler et al., 2011; Ward and Miwa, 1978). When male sperm are deposited into the uterus during mating, they activate and migrate to the spermatheca. In hermaphrodites, sperm are frequently pushed out of the spermatheca when a fertilized egg exits into the uterus. These sperm need to migrate back to the spermatheca. We previously observed that aged mutant hermaphrodites retained their sperm, suggesting that the sperm were motile and able to maintain their position in the spermatheca. We tested the motility of *spe-51(as39)* male sperm. We stained mated *fem-1(hc17)* mutants with DAPI to see whether the sperm made their way to the spermatheca. Both control and *spe-51(as39)* mutant sperm were able to migrate to the spermatheca (Fig.2G-I), suggesting that they activated properly *in vivo* and were motile. We conclude that *spe-51* is not required for sperm activation or sperm motility.

*C. elegans* male sperm can out compete hermaphrodite self-sperm at fertilization (LaMunyon and Ward, 1995; Singson et al., 1999; Ward and Carrel, 1979). A mated hermaphrodite preferably uses male sperm over self-sperm and as a result, the production of self-progeny is suppressed (LaMunyon and Ward, 1995). This effect can be measured by counting the number of outcross and self-progeny (Singson et al., 1999). We tested whether *spe-51(as39)* mutant sperm were also able to out compete hermaphrodite self-sperm. When males were given a 48-hour window to mate with the hermaphrodite, the production of self-progeny is suppressed with both control and *as39* males (Fig.2J, p<0.01 for both control and as39 males). This result suggests that *spe-51(as39)* males can out compete hermaphrodite self sperm despite lacking the ability to fertilize the egg.

### SPE-51 encodes a secreted protein with an Ig-like domain

Traditional mapping positioned the *as39* mutation on Chromosome IV between *dpy-13* and *unc-24* (data not shown). Using a mapping-by-sequencing approach (Doitsidou et al., 2010; Sarin et al., 2008; Thompson et al., 2015; Zhao et al., 2018), we identified a similar region as having a low frequency for Hawaiian SNPs (data not shown). We then searched for homozygous, nonsynonymous mutations that affect coding sequences within this region. We found a single nonsense mutation (Chr:IV 4696911CtoT) in the second exon of the gene *T22B11.1*. Previous gene expression profiling has shown that *T22B11.1* expression is enriched in males (Ortiz et al., 2014; Reinke et al., 2000). Sanger sequencing of *spe-51(as39)* mutants showed that this mutation indeed exists in the genome of the mutants. To confirm that this mutation caused the sterility phenotype in *spe-51(as39)*, we performed transgenic rescue experiments. A transgene carrying a wild-type copy of *T22B11.1* genomic DNA is able to rescue the fertility defect in the mutant (Fig. 1B). Thus, we confirm that *spe-51* is *T22B11.1*.

The *spe-51* gene produces a single transcript that encodes a protein of 468 amino acids. BLAST searches (https://blast.ncbi.nlm.nih.gov/Blast.cgi) showed no obvious homologs outside of nematodes. The nonsense mutation in the *as39* allele leads to a truncation of the protein at amino acid 31 (Q31STOP, Fig. 3A). We have since obtained two additional alleles of *spe-51*. The *as43* allele has an insertion that causes a frameshift that alters the coding sequence after amino acid 19 while the *as44* allele has a nonsense mutation that truncates the protein at amino acid 20 (Fig. 3A). Both *as43* and *as44* show the same sterility phenotype in hermaphrodites (Fig. 1B). Based on the nature of the molecular lesions in all three alleles and the lack of fertility at all culture conditions, they are all likely null alleles.

**Figure 3.**
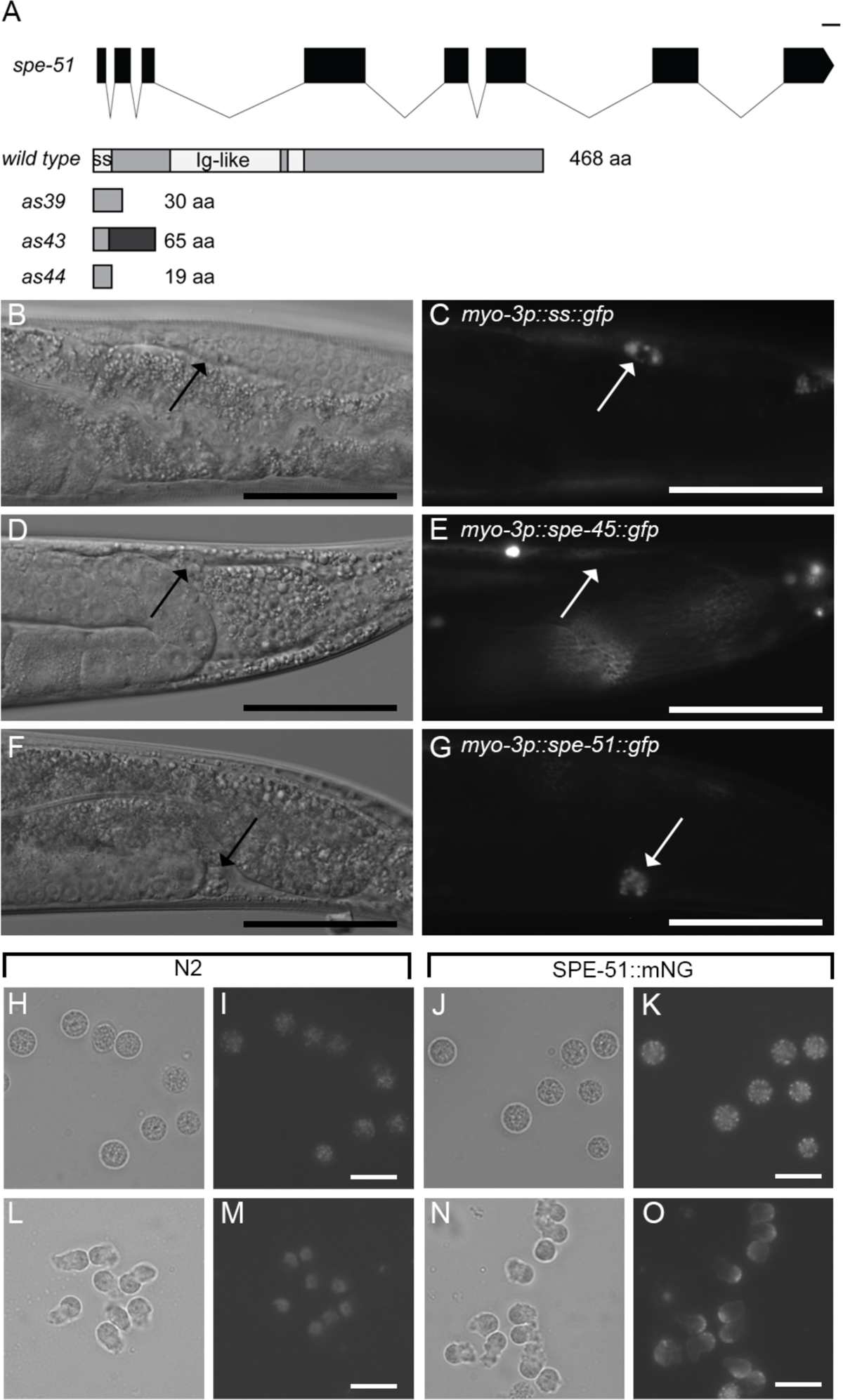
Molecular features, secretion, and localization of SPE-51. A, The *spe-51* gene and protein products from different alleles. Top panel shows the gene structure. Black boxes show the exons, and black lines in between exons show the introns. Bottom panels show the protein products. Wild-type SPE-51 protein: ss, signal sequence. The white box in the middle of the protein shows the hydrophobic region that some programs predict as a transmembrane domain. The mutant proteins predicted from all three mutant alleles are shown below the wild type. B-G, Expression of SPE-51::GFP in the muscle and its uptake by coelomocytes. B, D and F, DIC imaging. C, E and G, Green fluorescence of the worm as shown in A, C and E, respectively. Arrows in each panel point to the coelomocyte in the posterior region of the worm. Scale bars represent 50 μm. H-O, SPE-51 localization. H-K, spermatids. L-O, *in vivo* activated spermatozoa. Scale bars represent 10 μm.

The high similarity between *spe-51* phenotypes and those of the *spe-9* class mutants led us to predict that SPE-51 functions during sperm-egg interaction and must be a transmembrane protein just like all the other previously known members of the SPE-9 class proteins. SPE-51 is predicted to have a signal sequence at the N-terminus (SignalP). It is also predicted to have a single Ig-like domain (amino acid 76-190) (Fig. 3A) (Kelley et al., 2015). Although some programs (“DAS” and “Phyre2”) did recognize a transmembrane domain in SPE-51 (amino acids 197-212), other programs predicted that it is secreted (“TMHMM2.0” and “TOPCONS”). We decided to test whether it is secreted with a *C. elegans* secretion assay (Fares and Grant, 2002; Fares and Greenwald, 2001). Secreted proteins that are expressed from somatic tissues such as the gut or muscle are taken up by the scavenger cells, coelomocytes. We included two controls, a secreted version of GFP (ssGFP), and a SPE-45 tagged with GFP (Nishimura et al., 2015; Singaravelu et al., 2015). SPE-45 is a predicted transmembrane protein and is not expected to be secreted. We found that when expressed in the muscle, ssGFP is present in the coelomocytes, SPE-45::GFP is not (Fig. 3B-E). SPE-51::GFP is found in the coelomocytes indicating that it is secreted from the muscle cells (Fig. 3F-G). Similar results were seen when SPE-51::GFP was expressed in the gut (data not shown). We conclude that SPE-51 is not a transmembrane protein but is secreted.

It is intriguing that SPE-51 is a secreted protein. All previously characterized *spe-9* class genes encode transmembrane proteins (Krauchunas et al. 2016). Further, *spe-51* mutations impact sperm function at fertilization. If SPE-51 is indeed secreted by the sperm, we expect that the presence of SPE-51 secreted by wild-type sperm could allow the mutant sperm to gain the ability to fertilize oocytes. If *spe-51* mutant sperm were mixed with wild-type sperm, both mutant and wild-type sperm should be able to fertilize eggs. We re-analyzed our data from the male fertility test where we mated mutant males with either *fem-1* or *dpy-5* hermaphrodites. The *fem-1* mutant produces no sperm but normal oocytes whereas the *dpy-5* mutant produces sperm with functional SPE-51. During a 24-hour mating period, *spe-51(as39)* mutant males produced an average of 5.9(±1.2) cross progeny with *fem-1*, but 1.2(±0.3) outcross progeny with *dpy-5* (Fig.1C). These numbers are consistent with SPE-51 acting cell-autonomously. We further tested the question of autonomy by crossing SPE-51-positive males to mutant *spe-51(as39)* hermaphrodites, to rule out the contribution of seminal fluid. For this experiment, we used *spe-9(eb19)* mutant males that are completely sterile so any progeny can be counted as self-progeny from *spe-51(as39)* hermaphrodites. As a control, we used *spe-51(as39)* males.

Compared to unmated hermaphrodites, hermaphrodites mated with either *spe-9(eb19)* or *spe-51(as39)* males did not show an increase or decrease in the number of self-progeny (Table S1). These data argue that SPE-51 acts cell-autonomously and stays associated with the sperm that produced the protein.

### SPE-51 localization

To visualize the SPE-51 protein, we obtained a tagged allele of *spe-51* by inserting a C-terminal mNeonGreen (mNG) tag at the endogenous location using Crispr. In whole-mount males, we detected a signal in the region where spermatids are stored (data not shown). This observation suggests that SPE-51 is secreted by the sperm. In spermatids, we detected a signal within the sperm that is consistently above the background autofluorescence from control spermatids (Fig. 3H-K). The pattern of signal is consistent with localization to the membranous organelles (MOs) (Fig. 3K). In *in vivo* activated spermatozoa, the signal decorated the whole sperm surface, including the pseudopods (Fig. 3L-O). In most of the spermatozoa, the signal can also be seen in what appears to be the MOs, which have fused with the plasma membrane during spermiogenesis. These results indicate that SPE-51 is secreted from the MOs and stays associated with the mature sperm. Further, this localization pattern is consistent with the cell-autonomous behavior.

## Discussion

### SPE-51’s functions and conserved roles of Ig domains during fertilization

Here, we show that the *spe-51* gene is dispensable for early spermatogenesis and sperm activation but is required for sperm function at fertilization. *spe-51* mutant animals phenocopy mutants of the *spe-9* class genes that encode proteins that are thought to form the fertilization synapse at the interface between the sperm and the oocyte (Krauchunas et al., 2016; Singson, 2001). The mutant phenotype of *spe-51* establishes it as a new member of the *spe-9* class of sperm function genes and as a unique component of the fertilization synapse. In another manuscript by Krauchunas et al., we also show that *spe-36* is a secreted protein required for fertilization in the sperm. Together, this is the first time that sperm secreted molecules are shown to be involved in the interactions between the sperm and egg with genetic validation for function.

Ig superfamily molecules are generally implicated in the molecular recognition and binding activities of cells (Barclay, 2003). The complementarity-determining region of the Ig-domain is thought to mediate specificity of ligand binding. If the Ig domain (or other parts) of SPE-51 functions to bind other proteins, it will be important to determine if they are *cis* binding partners on the surface of sperm, *trans* binding partners on the surface of the egg, or both (Fig. 4). These interactions may serve as an initial recognition signal or trigger functional conformational changes in SPE-51 itself and/or its binding partners on either gamete surface. Interestingly, the Ig domain of IZUMO1 is not required for interactions with JUNO (Aydin et al., 2016; Nishimura et al., 2016; Ohto et al., 2016), raising the possibility that IZUMO1 may interact with additional unknown molecules in the fertilization synapse. For both SPE-51 and IZUMO1, these interactions and molecular dynamics can further organize or regulate the biochemical activity of other components of the fertilization synapse.

**Figure 4.**
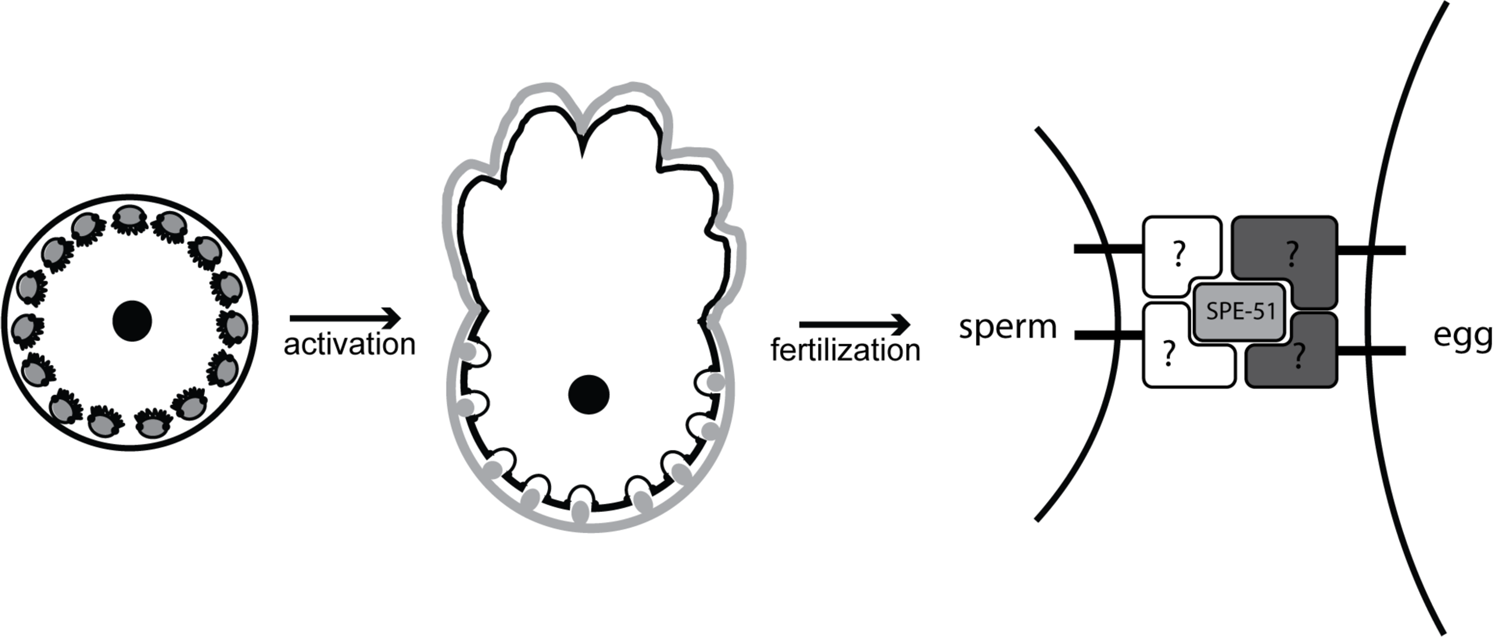
Model of SPE-51 localization and how it functions within the fertilization synapse. We predict that spermatids (shown on the left) express SPE-51 in the MOs (shown as light grey shapes within the cytoplasm of spermatids). Upon activation, SPE-51 is translocated to the extracellular space of spermatozoa but stays associated with them (middle panel) possibly via interactions with unknown proteins on the sperm side. Upon fertilization when the sperm encounters the oocyte, SPE-51 may also interact with proteins (dark grey) on the oocyte surface.

Several Ig superfamily molecules have been shown to play a role in fertilization in both *C. elegans* and mammals, including mammalian IZUMO1 (Inoue et al., 2005), SPACA6 (Barbaux et al., 2020; Lorenzetti et al., 2014; Noda et al., 2020), and SPE-45 in worms (Nishimura et al., 2015; Singaravelu et al., 2015). Disruptions of both *Izumo1* and *Spaca6* result in male sterility with mutant spermatozoa being able to migrate into the oviduct, to undergo acrosome reaction and to penetrate the zona pellucida but unable to fertilize the egg (Inoue et al., 2005; Lorenzetti et al., 2014). These phenotypes strongly resemble those of the *spe-45* or *spe-51* mutants, suggesting that Ig superfamily proteins may have an evolutionarily conserved role during fertilization (Nishimura and L’Hernault, 2016; Tajima and Nishimura, 2018). In fact, a chimeric SPE-45 protein with its Ig domain replaced by that of IZUMO1 partially rescues mutant *spe-45* sterility (Nishimura et al., 2015). These data further suggest a deep structural conservation at the fertilization synapse in evolutionary distant species.

The SPE-51 protein contains a stretch of hydrophobic amino acids (aa197-212). If this region is not functioning as a transmembrane domain, what does it do? Hydrophobic regions of molecules can be used in protein interactions (Tsai et al., 1997; Young et al., 1994). More intriguingly, hydrophobic regions can function as fusion peptides as seen in viral fusion proteins (Brukman et al., 2019; Hernandez and Podbilewicz, 2017; Perez-Vargas et al., 2021). Known gamete fusogens such as HAP2 are transmembrane proteins (Fedry et al., 2017; Valansi et al., 2017) whereas SPE-51 is secreted. If the internal hydrophobic domain of SPE-51 does function as a membrane fusion peptide during fertilization, it will likely require some other molecule(s) to bind to it and to bring it close to its target membrane in either sperm or eggs. Without these theoretical binding partners, it would be difficult to imagine that SPE-51 could function by itself in established cell-cell fusion assays (Hernandez and Podbilewicz, 2017). SPE-51 would be necessary but not sufficient to drive sperm-egg fusion in worms.

### Secretion and tethering of SPE-51

Regardless of the precise biochemical function of SPE-51, an important question is how the SPE-51 protein is secreted. First, we think that SPE-51 is unlikely secreted by cell types other than the sperm. The gene is expressed and enriched in the male germline (Ortiz et al., 2014; Reinke et al., 2000). This expression together with the sperm-specific phenotype, cell autonomous behavior, and sperm-associated localization patterns supports the idea that the sperm express and secrete SPE-51. Proteins in the sperm could take a few paths to secretion. During sperm activation, intracellular Golgi-derived vesicles, called the MOs, fuse with the plasma membrane (Ward et al., 1981), in a process that is analogous to the acrosome reaction in sperm from other species (Nishimura and L’Hernault, 2010). This fusion leads to the release of the contents of the MO extracellularly and the translocation of molecules onto the plasma membrane (Ward et al., 1983). One possibility is that SPE-51 is packed into the MOs in spermatids and released to be associated with the plasma membrane upon MO fusion. The fact that SPE-51::mNG can be seen in what appears to be MOs in spermatids and fused MOs in the spermatozoa suggests this is likely the case. Two other sperm proteins required for fertilization, SPE-38 and SPE-41, co-localize to the MOs in spermatids and during spermiogenesis translocate to the pseudopod or the whole plasma membrane, respectively (Chatterjee et al., 2005; Singaravelu et al., 2012; Takayama and Onami, 2016). Alternatively, SPE-51 may be secreted via classical secretory pathways, by either spermatids or spermatozoa, similar to the secretion of Major Sperm Proteins (Miller et al., 2001). We think this secretion pathway is unlikely due to our detection of SPE-51::mNG in the MOs. After secretion the protein then stays associated with the sperm by interactions with other proteins or by post-translational modifications.

While we cannot rule out lipid modifications as tethers, it is likely that some sperm-surface proteins tether SPE-51 to the sperm surface. One such candidate is SPE-45, given that Ig domains most frequently interact with other Ig domains (Barclay, 2003). Another candidate is the four-pass transmembrane protein SPE-38. Although SPE-38 is thought to localize to the pseudopods of spermatozoa, it is required for the proper distribution of SPE-41 on the entire spermatozoa surface (Singaravelu et al., 2012) and interacts with multiple sperm surface proteins in our previous *C. elegans* sperm interactome studies (Marcello et al., 2018). Thus, it is possible that SPE-38 facilitates the secretion and distribution of SPE-51 as well. Future work on identifying these possible interactors will point to the mechanisms of the assembly and dynamics of the fertilization synapse.

### Secreted proteins in modulating gamete functions

Proteins secreted by the male and female reproductive tract and by the sperm acrosome modify the sperm surface to promote sperm maturation, capacitation, and function (Bjorkgren and Sipila, 2019; Fujihara et al., 2019; Hirohashi and Yanagimachi, 2018; Kiyozumi et al., 2020; Suarez, 2008, 2016). In *C. elegans*, secreted proteins modify the sperm to modulate its differentiation and function. For example, the secreted protease inhibitor SWM-1 puts the secreted male sperm activator TRY-5 in check to ensure proper timing of sperm activation (Chavez et al., 2018; Smith and Stanfield, 2011; Stanfield and Villeneuve, 2006). Our observations that SPE-51 is a secreted protein required for fertilization suggest that secreted proteins on the sperm surface can affect the ability of the sperm to interact with the egg (see Krauchunas et al. on SPE-36 on Biorxiv). These observations together provide evidence that the sperm surface is modified by factors secreted by both the sperm itself and by the environment, and that these modifications serve as a necessary step to ensure fertilization competency.

Here we have shown that the *spe-51* gene in *C. elegans* functions during gamete interactions. To our knowledge, this is the first example in any model organism that a secreted protein is required for the sperm to interact with and fertilize the egg (see Krauchunas et al. on SPE-36 on Biorxiv). Our discovery serves as a paradigm for understanding how secreted proteins modify the sperm surface to influence their ability to interact with the egg. The presence of Ig-like domains in worm SPE-45 and SPE-51 and in mammalian IZUMO1 and SPACA6 highlights important functions of Ig-domains in gamete interactions during fertilization. These findings provide insights into the complexity of the fertilization synapse and guide our understanding about modifications to the sperm that affect their fertilization competency.

## Methods

### Strain maintenance

All the strains are maintained as described previously (Brenner, 1974). The following strains are used in this study: wild type N2 Bristol, Hawaiian CB4856, *him-5(e1490)*, *fem-1(hc17)*, *dpy-5(e61)*, arIs37, *spe-51(as39)*, *spe-51(as43)*, *spe-51(as44),* and *spe-9(eb19)*. The *spe-51(as39)* strain was identified in a previous ethyl methanesulfonate (EMS) mutagenesis screen for fertility genes by Dr. Guna Singaravelu and Dr. Diane Shakes in the lab of Dr. Andrew Singson. Details of the screen was previously described in (Singaravelu et al., 2015). Before transgenic rescue, *spe-51(as39)* was maintained by crossing to N2 males and selecting sterile worms in the F2 generation. Both *spe-51(as43)* and *spe-51(as44)* were isolated by Crispr-mediate genome editing tools. We used synthetic sgRNA purchased from Synthego and followed the *dpy-10* co-conversion strategy describe by Paix et al (Paix et al., 2017). The strain PHX2737, *spe-51(syb2737)* carries a mNeonGreen insertion at the endogenous location on the C-terminus. This allele was generated by SunyBiotech.

### Light Microscopy

Hermaphrodites were mounted onto agarose pads and paralyzed in 1mM levamisole. To image the sperm, hermaphrodites or males were dissected in sperm media on a glass slide. DIC imaging was done using a Zeiss Universal microscope and captured with a ProgRes camera with the ProgresCapturePro software.

### DAPI staining

Hermaphrodites were fixed in cold methanol for 30 seconds and washed in PBS. They were then transferred onto agarose pads with Vectashield mounting medium with DAPI (Vector Laboratories).

### Progeny and ovulation count

For hermaphrodite fertility, each hermaphrodite was picked onto a fresh plate at the L4 stage. They were then transferred daily onto a new plate until they stopped laying eggs (for progeny count) or oocytes (for ovulation count). The number of adult progeny or progeny plus oocytes was counted daily.

To assess male fertility, *fem-1(hc17)* hermaphrodites were crossed with *him-5(e1490)* or *spe-51(as39); him-5* young adult males in a 1:4 ratio. After 24 hours, males were removed and hermaphrodites were transferred to a new plate. The hermaphrodites were transferred daily until they stopped laying eggs. The number of adult progeny was counted. Hermaphrodites that died or became absent before the end of the experiment were excluded from the analyses.

### Sperm activation

L4 stage males were isolated from hermaphrodites overnight. The next day, they were dissected in sperm media with or without Pronase (200 μg/mL) (Shakes and Ward, 1989). Imaging was done 10 minutes after dissection.

L4 stage hermaphrodites were isolated from males for overnight. Then their reproductive tract was dissected in sperm media without Pronase.

### Sperm migration

*fem-1(hc17)* hermaphrodites were crossed with *him-5(e1490*) or *spe-51(as39); him-5(e1490)* young adult males in a 1:4 ratio. After 24 hours, hermaphrodites were used for DAPI staining.

### Sperm competition

Individual *dpy-5(e61)* hermaphrodites were crossed with *him-5(e1490*) or *spe-51(as39); him-5(e1490)* young adult males in a 1:4 ratio for 48 hours at 20°C. At the end of 48 hours, the parents were removed. Three days later, all the Dpy and non-Dpy progeny on the plates were counted.

### Sperm mixing to test cell autonomy

Individual *spe-51(as39)* hermaphrodites were crossed with *spe-9(eb19); him-5(e1490)* males, or *spe-51(as39); him-5(e1490)* males in 1:3 ratio for 24 hours at 20°C. At the end of 24 hours, the parents were removed. Three days later, all progeny on the plates were counted.

### Whole genome sequencing and data analyses

Homozygous *spe-51(as39)* was crossed with CB4856 males. F1 progeny were allowed to self-fertilize. About 500 F2 progeny were picked into individual wells of 24-well plates at the L4 stage. Their fertility phenotype was scored the next day. About 100 sterile F2 worms were identified and pooled for genomic DNA prep by standard column purification (Qiagen Minelute PCR purification Kits).

Library construction was done the same way as described previously (Krauchunas et al., 2018). For data analysis, we used the MiModD program and filtered the region from 4.5Mbp∼7Mbp on Chromosome IV.

### Transgenic rescues

The genomic region of *spe-51* was amplified by PCR from N2 genomic DNA using the Expand Long Template PCR system from Roche (Catalog Number 11681834001). The following primers were used: genomic_Fwd: 5’-TTTATGCCTGCCCGCCTATG-3’, genomic_Rev: 5’-TCCTCGAAATCGGCTGAAATGA-3’. This 6.5 kb product covers all the exons as well as 1378 bp upstream of the transcription start site and 1130 bp downstream of the stop codon. The PCR product (2uL of 120ng/μL) together with the injection marker plasmid myo-3p::gfp (3μL of 180 ng/μL) was injected into N2 young adult worms and stable transgenic lines were selected. This transgenic line was then crossed into the *spe-51(as39)* mutant background. F2 individuals that segregated GFP-negative sterile progeny indicated that they were homozygous for *spe-51(as39).* GFP-positive worms from these individuals were then scored for fertility to evaluate rescue.

### Secretion assay

The plasmid pJF25 was used as a backbone for cloning. This plasmid contains the promoter region of *myo-3*, a signal sequence from SEL-1, GFP, and the 3’UTR of *unc-54*. For cloning, pJF25 was digested by AgeI and XbalI which flank the signal sequence from SEL-1. Digested DNA was purified by column purification (Qiagen PCR purification Kit). Gene fragments for *spe-51* (cDNA) and *spe-45* (genomic DNA) were amplified by PCR, purified by column, and inserted into digested pJF25 by Gibson Assembly (NEBuilder HiFi DNA assembly kit). We designed the Gibson Assembly so that *spe-51* or *spe-45* would be inserted between the *myo-3* promoter and GFP, replacing the original signal sequence. For expression in the gut, ssGFP and SPE-51::GFP were amplified from the above plasmids. All three genes were then inserted to a plasmid containing the *vit-2* promoter by Gibson assembly. All the final constructs were verified by Sanger sequencing. For injections, individual expression constructs were mixed with pCFJ90 and injected into the gonad of young adults. F1 transformants were isolated and F2 transmitting lines were generated. At least two independent lines for each expression construct were obtained and imaged.

### Localization of SPE-51

The strain PHX2737 was heat-shocked to obtain males. It was then maintained by continuously mating the males with PHX2737 hermaphrodites. For spermatids, males were separated from hermaphrodites overnight and then dissected in sperm media. To get *in vivo* activated sperm, 10 hermaphrodites and 20 males were picked on to a plate and allowed to mate overnight. The next day, hermaphrodites were dissected in sperm media. Following dissection, sperm were then dispersed on the slide and ready for imaging. N2 crosses were set up the same way as a control.

Images were obtained on a AxioPlanII epifluorescent microscope with an ORCA charge-coupled device camera (Hamamatsu) using the iVision software (Biovision Technologies).

## Acknowledgements

We thank Dr. Christopher Rongo for sharing experimental equipment. We thank Dr. Matthew Marcello and all members of the Singson lab for critical feedback on this manuscript. Research in the Singson lab is supported by a grant from the NIH to A.W.S (R01HD054681). X.M. was supported by a Busch Postdoctoral Fellowship.

## Author contributions

Conceptualization, X.M., A.R.K., A.W.S.; Methodology, X.M., A.R.K., B.D.G.; Investigation, X.M., M.D., S.D., J.N., G.S., S.G.G.; D.C.S.; Writing - Original Draft, X.M., A.W.S.; Writing - Review & Editing, X.M., A.R.K., A.W.S.; Supervision, A.W.S.; Funding Acquisition, A.W.S.;

---

**Figure S1.**
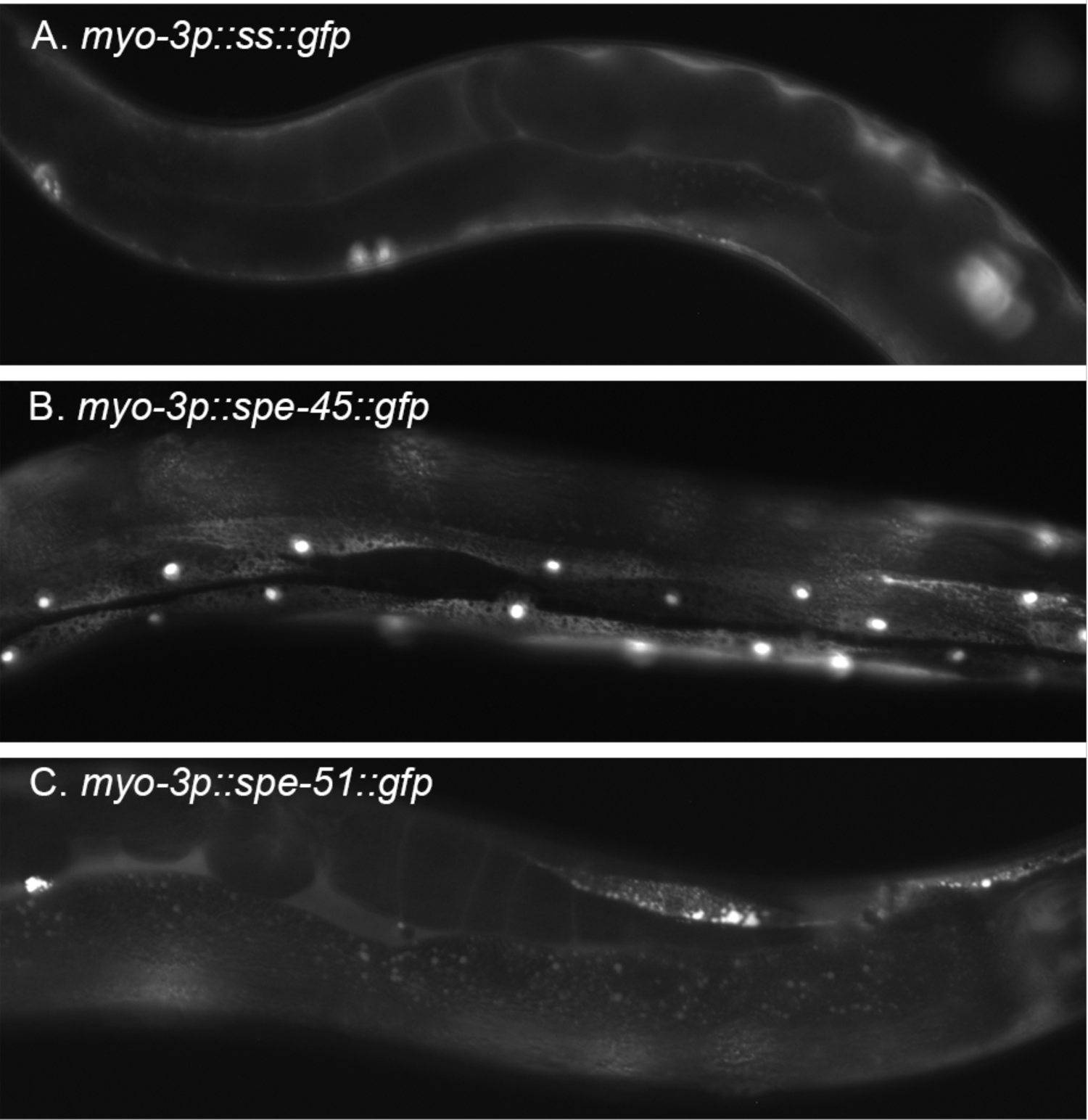
Expression of SPE-45::GFP and SPE-51::GFP in the muscle.

**Table S1.** SPE-51 acts cell autonomously. Progeny counts of mated and unmated *spe-51(as39)* hermaphrodites, shown as mean±SEM. ns, not statistically significant.

| Males | # of cross progeny |
| --- | --- |
| N/A | 0.43±0.11 |
| *spe-9(eb19)* | 0.32±0.13 (ns) |
| *spe-51(as39)* | 0.67±0.21 (ns) |

